# Tau Signaling in Injured Spinal Cord

**DOI:** 10.64898/2026.09.19.752874

**Authors:** Natasha Bukreska, Sienna R. Casciato, Tatiana Bezdudnaya

## Abstract

Tau is a microtubule-associated protein important for neuronal structure and function, with abnormal phosphorylation associated with neuronal dysfunction and degeneration. While tau pathology has been extensively studied in neurodegenerative diseases and traumatic brain injury, its role in spinal cord injury (SCI) remains unclear. This study investigates changes in the levels of tau isoforms and phosphorylation following a high lateral C2 hemisection (C2Hx) in adult female rats. Western blot analysis was performed on spinal cord tissue collected caudal to the lesion at 4 weeks post-injury. Low-molecular-weight (LMW) tau and high-molecular-weight, Big tau did not differ significantly between injured and control groups. In contrast, phosphorylated tau (p-tau), detected using the AT8 antibody, was significantly elevated after injury, suggesting a shift in tau functional state. Assessment of microtubule stability markers revealed increased acetylated tubulin and decreased tyrosinated tubulin after SCI, indicating a shift toward a more stable cytoskeletal state. Immunohistochemistry was performed on spinal cord sections at 6 weeks post-injury and dorsal root ganglia (DRG) at 8 weeks post-injury. These analyses demonstrated increased p-tau immunoreactivity within the ipsilateral dorsal horn of the injured spinal cord and a higher proportion of p-tau-positive neurons in DRGs from injured animals compared with controls. Together, these findings demonstrate that cervical SCI induces persistent tau phosphorylation without altering total tau levels, and that this modification extends beyond the lesion site into distal spinal regions and beyond the acute phase of injury. These alterations may influence neural plasticity and remodeling in the injured spinal cord.

## 1. Introduction

Tau proteins are microtubule-associated proteins highly expressed in neurons, where they regulate microtubule organization and cytoskeletal dynamics (Baas & Qiang, 2019). As a major component of the axonal cytoskeleton, tau maintains axonal integrity and supports intraneuronal transport, making proper tau regulation essential for neuronal function (Wang & Mandelkow, 2016). The MAPT gene produces multiple tau isoforms through alternative mRNA splicing (Wiche et al., 1991; Fischer & Baas, 2026). These isoforms are broadly categorized into low-molecular-weight (LMW) tau and high-molecular-weight, Big tau. LMW tau consists of six isoforms predominantly expressed in the central nervous system (CNS) (Corsi et al., 2022; Fischer, 2023; Fischer & Baas, 2026), whereas Big tau contains a large exon 4a-encoded projection domain that distinguishes it structurally from LMW tau.

Tau isoform expression varies across neuronal populations and developmental stages. While LMW tau is widely expressed, Big tau is enriched in neurons with exceptionally long axons and high transport demands, including dorsal root ganglia (DRG), sympathetic ganglia, spinal motor neurons, and selected CNS regions (Fischer & Baas, 2026; Jin et al., 2023; Boyne et al., 1995). The extended projection domain of Big tau increases microtubule spacing and may reduce pathological tau aggregation, potentially providing protection against tau-related degeneration (Fischer & Baas, 2026). However, these structural adaptations may also influence microtubule dynamics and neuronal plasticity, suggesting that tau isoforms may contribute differently to neuronal responses following injury.

Tau phosphorylation is a critical post-translational modification that regulates tau function. Under physiological conditions, reversible and site-specific phosphorylation controls tau interactions with microtubules and contributes to regulation of axonal transport (Trushina et al., 2019). Following injury, disruption of kinase and phosphatase balance can result in tau hyperphosphorylation, causing tau detachment from microtubules and affecting cytoskeletal stability (Hung et al., 2005; Yu et al., 2005). Chronic dysregulation of tau phosphorylation contributes to tauopathies, including Alzheimer’s disease and frontotemporal dementia with Parkinsonism-17, where hyperphosphorylated tau forms abnormal aggregates associated with neuronal dysfunction (Lee et al., 2001; Avila, 2006).

Spinal cord injury (SCI) causes extensive neural dysfunction beyond the primary lesion site, resulting in persistent sensory and motor impairments (Kim et al., 2025). Because tau is closely associated with axonal microtubules and cytoskeletal organization, it has emerged as a potential biomarker of axonal injury following SCI (Caprelli et al., 2018). Previous studies have demonstrated an acute increase in phosphorylated tau (p-tau) following SCI, with elevated p-tau immunoreactivity observed at the injury epicenter and extending into surrounding spinal regions. P-tau-positive axons peaked at 1-day post-injury and progressively declined through 7 days (Caprelli et al., 2018). Additionally, increased tau levels correlate with injury severity and poorer motor recovery, while inhibition of tau phosphorylation may reduce neuroinflammation and oxidative stress and improve functional outcomes following SCI (Tang et al., 2019; Chen et al., 2023).

Despite these findings, important gaps remain regarding the long-term regulation of tau following SCI. Previous studies have primarily focused on acute changes in total tau or localized injury responses, while the effects of SCI on specific tau isoforms, chronic tau phosphorylation, and distal spinal regions remain unclear. We hypothesize that SCI induces persistent tau phosphorylation associated with altered microtubule stability without altering total tau abundance, and that these changes extend beyond the lesion site into distal spinal regions.

## 2. Materials and methods

### Animals

Adult female Sprague-Dawley rats (250-275 g; Envigo) were used in this study (n=18). All experiments were performed with approval from the Institutional Animal Care and Use Committee at Drexel University (IACUC protocols LA-23-728 and 20923). At the experimental endpoint, all animals were euthanized by an overdose of Euthasol. Animals were either transcardially perfused for immunohistochemical analyses or rapidly dissected to collect fresh spinal cord tissue for Western blot analysis.

### Injury model

The present study used a lateral C2 hemisection (C2Hx) injury model immediately rostral to the phrenic motoneuron pool, which is widely used to investigate respiratory impairments following spinal cord injury (SCI) (Hoh et al., 2013). This model directly interrupts supraspinal input to ipsilateral phrenic motoneurons and results in ipsilateral hemidiaphragm paralysis (Figure 1). The C2Hx model was selected because it produces a reproducible unilateral disruption of descending pathways, including bulbospinal inputs to the phrenic motor nucleus, allowing assessment of injury-induced changes within remaining respiratory circuits.

**Figure 1.**
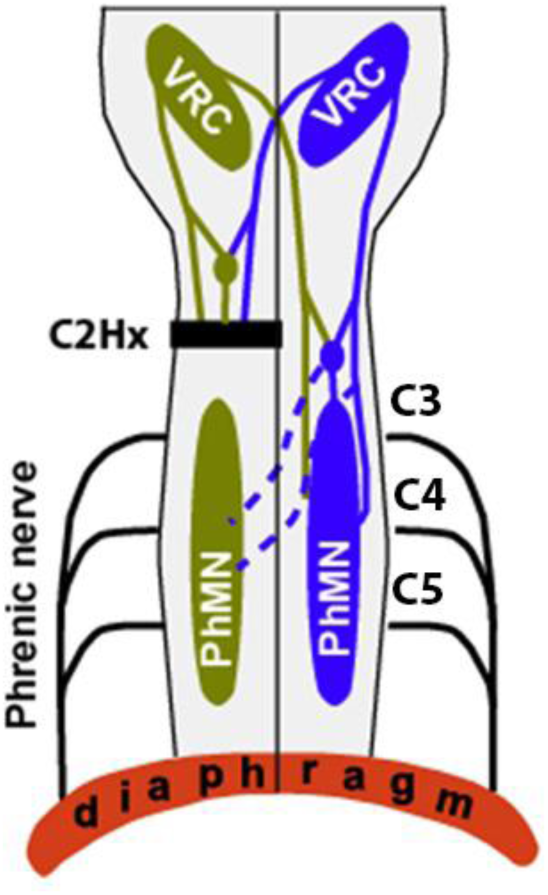
Schematic diagram of the left (brown) and right (blue) phrenic circuitry showing the location of C2Hx injury. The ventral respiratory column (VRC) in the brainstem projects directly onto spinal phrenic motoneurons (PhMNs), and spinal pre-motor interneurons, on each side of the spinal cord. PhMNs innervate the ipsilateral hemi-diaphragm via a single phrenic nerve. Note that some VRC projections decussate at the brainstem or spinal levels.

In addition to spinal cord analyses following C2Hx, dorsal root ganglia (DRG) samples provided by Dr. Detloff’s laboratory were included in this study. These tissues were obtained from animals subjected to a unilateral C5 contusion injury using an IH impactor (Precision Systems and Instrumentation, Lexington, KY, USA) at the force of 200 kdyn. Ipsilateral DRGs from the C6–C7 levels were collected at 8 weeks post-injury, representing the chronic phase of the response. The dataset included samples from control (n = 2) and injured (n = 2) animals.

### Spinal cord injury surgery

All surgeries were performed under ketamine (80–120 mg/kg, i.p.) and xylazine (20 mg/kg, s.c.) anesthesia. After achieving a surgical plane of anesthesia, the dorsal neck region was shaved and aseptically prepared. A midline incision was made from the base of the skull to the level of the shoulder blades, extending through the skin and overlying musculature to expose the cervical vertebrae. A partial laminectomy was performed at the C2 vertebral level.

The dura mater was incised (∼5 mm) rostrocaudally along the midline of the dorsal spinal cord. A C2Hx was then performed immediately caudal to the C2 rootlets, extending from the midline to the lateral edge of the spinal cord using a No. 11 scalpel blade. This procedure produced a unilateral cervical spinal cord injury by disrupting the ipsilateral descending pathways while preserving contralateral circuitry.

Following the lesion, the surrounding muscles were sutured (4–0 sterile suture), and the skin was closed with wound clips. Postoperatively, Antisedan (0.2 mg/kg, s.c.) was administered to reverse xylazine. Lactated Ringer’s solution (5 mL, s.c.) and buprenorphine (0.03 mg/kg, s.c.) were administered for fluid support and analgesia. Animals received standard postoperative monitoring, including additional lactated Ringer’s injections, Nutri-Cal nutritional supplementation, routine cleaning, and daily body weight monitoring. Surgical procedure for unilateral C5 contusion is described here (Giddings & Detloff, 2026).

### Western blot

Following euthanasia, cervical spinal cord tissue caudal to the injury site (C3–C8) was rapidly dissected, immediately frozen on dry ice to preserve protein integrity, and subsequently divided into ipsilateral (left) and contralateral (right) halves. Only the ipsilateral spinal cord tissue was used for Western blot analysis. Fresh spinal cord and DRG tissues were homogenized in cold lysis buffer (RIPA: 25 mM Tris-HCl, pH 7.6; 150 mM NaCl; 1% NP-40 or Triton X-100; 1% sodium deoxycholate; 0.1% SDS) with a protease inhibitor cocktail using a Fisherbrand Sonic Dismembrator. Protein concentration was determined using the Pierce™ BCA Protein Assay Kit (Thermo Fisher Scientific, Waltham, MA, USA). Protein samples were diluted in 4× loading buffer (final 1×) with 2-mercaptoethanol and heated at 95°C for 5 min. Samples, together with 2 µL of Chameleon Duo pre-stained protein ladder (LI-COR, Lincoln, NE, USA), were separated by SDS-PAGE on 4–12% Bis-Tris polyacrylamide gels (Invitrogen, Waltham, MA, USA). Proteins were transferred onto PVDF membranes using a dry transfer system (Thermo Fisher iBlot, Waltham, MA, USA). Membranes were washed in Tris-buffered saline (TBS, pH 7.4) and blocked in Intercept TBS blocking buffer (LI-COR) for 1 h at room temperature. Membranes were then incubated overnight at 4°C with primary antibodies against Big tau (1:1000) (Jin et al., 2023), 3′ tau (1:5000) (Jin et al., 2023), phosphorylated tau (Ser202, Thr205, monoclonal antibody AT8, Thermo Fisher Scientific, 1:1000), acetylated tubulin (Millipore, T6793-2ML, 1:2000), GAPDH (1:14000), tyrosinated tubulin (mouse monoclonal tyrosine tubulin, Sigma, T9028, 1:1000), cofilin (1:14000), and actin (Millipore, Burlington, MA, USA; 1:70,000).

After washing in TBS, membranes were incubated with IRDye-conjugated secondary antibodies (LI-COR, 1:50,000) for 1 h at room temperature. Following several washes in TBS and a final rinse in distilled water, fluorescent signals were detected at 700 and/or 800 nm using the Odyssey® CLx Imaging System (LI-COR). All antibodies were diluted in Intercept Antibody Diluent (TBS).

### Immunohistochemistry

After completion of the experiments, rats were euthanized by overdose of Euthasol (0.5 mL, i.p.) and transcardially perfused with 0.9% NaCl solution followed by 4% paraformaldehyde (PFA). The spinal cords were carefully dissected and post-fixed in 4% PFA at 4°C overnight. Tissues were then blocked and cryoprotected subsequently in 15% and 30% sucrose solutions prepared in 0.1 M phosphate-buffered saline (PBS). Spinal cords were sectioned at 20 µm using a cryostat and collected in five series, such that each slide contained every fifth section, allowing for up to five different immunostainings. DRG samples were sectioned at 10 µm.

First, sections were washed in PBS (3 × 5 min), blocked against endogenous peroxidase activity (30% methanol, 0.6% hydrogen peroxide in PBS, incubated for 1 h), re-washed in PBS, and blocked against non-specific protein staining (10% donkey serum in PBS with 0.3% Triton-X, incubated for 1 h). Sections were then incubated overnight at 4°C with the following primary antibodies: rabbit anti-Big Tau (1:1000) (Jin et al., 2023), rabbit anti-3′tau (1:1000) (Jin et al., 2023), and mouse anti-p-tau (AT8) (1:500).

The following day, sections were washed in PBS (3 × 5 min) and incubated with secondary antibodies conjugated to different Alexa Fluor dye molecules for 2 h at room temperature. The following secondary antibodies were used: donkey anti-rabbit Alexa Fluor 488 (1:400, Jackson ImmunoResearch Laboratories Inc., #711-545-152) and donkey anti-mouse Alexa Fluor 647 (1:800, Invitrogen, #A31571). Stained sections were washed and coverslipped with Fluoromount-G mounting medium (Invitrogen, Thermo Fisher Scientific).

### Data analysis

All slides were examined, and images were acquired using a Leica Thunder microscope. Fluorescent staining was quantified using ImageJ (version 1.54g), and Western blot analysis was performed using Image Studio Lite (version 5.2).

For immunohistochemical analyses, measurements from multiple sections from the same animal were averaged to obtain a single value for each animal, with animals treated as independent biological replicates. Statistical analyses were performed using OriginPro 2025. Comparisons between control and injured groups were performed using independent-samples t-tests. Statistical significance was defined as p < 0.05. Because only two animals per group were available for the DRG analysis, these data were presented descriptively and were not subjected to statistical analysis.

## 3. Results

### Quantification of injury-induced changes in tau isoform levels and tau phosphorylation in the spinal cord

To evaluate the long-term molecular response to spinal cord injury, tissue samples collected caudal to the C2Hx lesion were analyzed at 4 weeks post-injury using Western blots. This time point represents an early chronic phase following injury. Western blot analysis was performed on ipsilateral cervical tissue to determine if the 4-week recovery period altered the protein level of specific tau isoforms. Western blot analysis of tau isoforms is shown in Figure 2. Figure 2A presents representative blots obtained using a 3′ tau antibody which detects all tau isoforms, while Figure 2B shows blots probed with a Big tau antibody, confirming the presence of the high molecular weight tau band (∼110 kDa) observed in Figure 2A. Actin (∼42 kDa) was used as a loading control.

**Figure 2.**
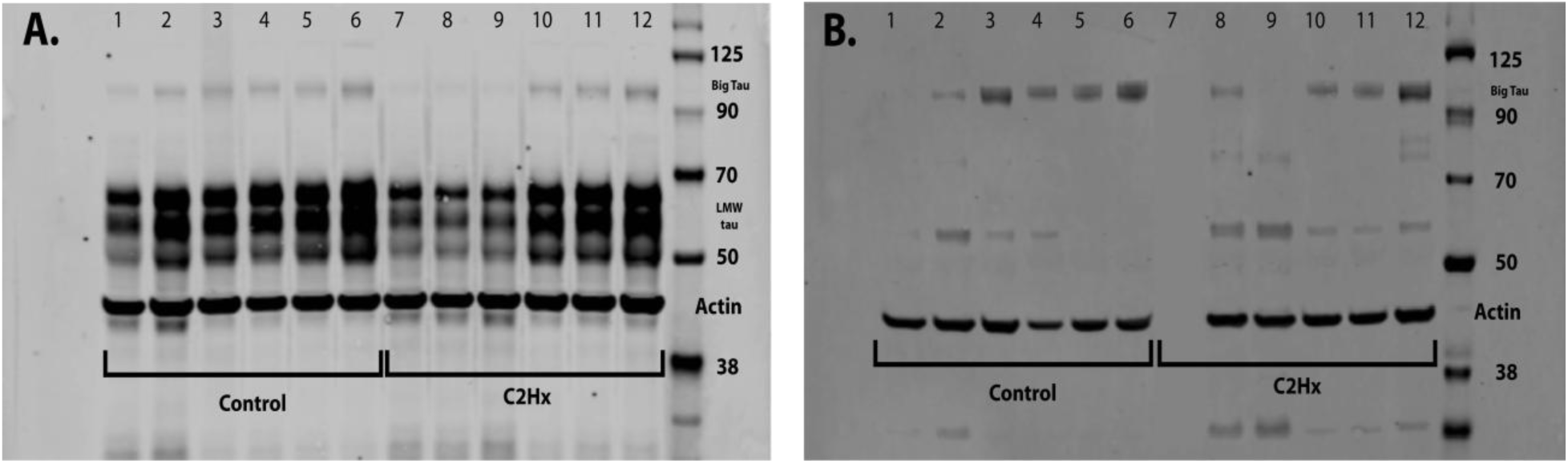
Representative Western blots showing tau isoform expression. (A) 3′ tau antibody detects LMW tau isoforms (∼50–65 kDa) and Big Tau isoform (∼110 kDa). (B) Big tau antibody detects high molecular weight tau (∼110 kDa). Actin (∼42 kDa) was used as a loading control. Lanes 1–6 represent control (CTR) animals, and lanes 7–12 represent C2Hx (injured) animals. Note, lane 7 is empty in B.

Quantification of LMW tau and Big tau, normalized to Actin, is shown in Figure 3. No statistically significant differences were observed between control and injured groups (LMW tau: p = 0.05; Big tau: p = 0.3, ratio Big Tau/LMW tau p = 0.7; T-test; n = 6 per group). However, a trend toward decreased expression of both LMW and Big tau was observed in injured animals. These results suggest that the overall pool of tau isoforms remains relatively stable within the post-lesional cervical spinal cord at 4 weeks post-injury.

**Figure 3.**
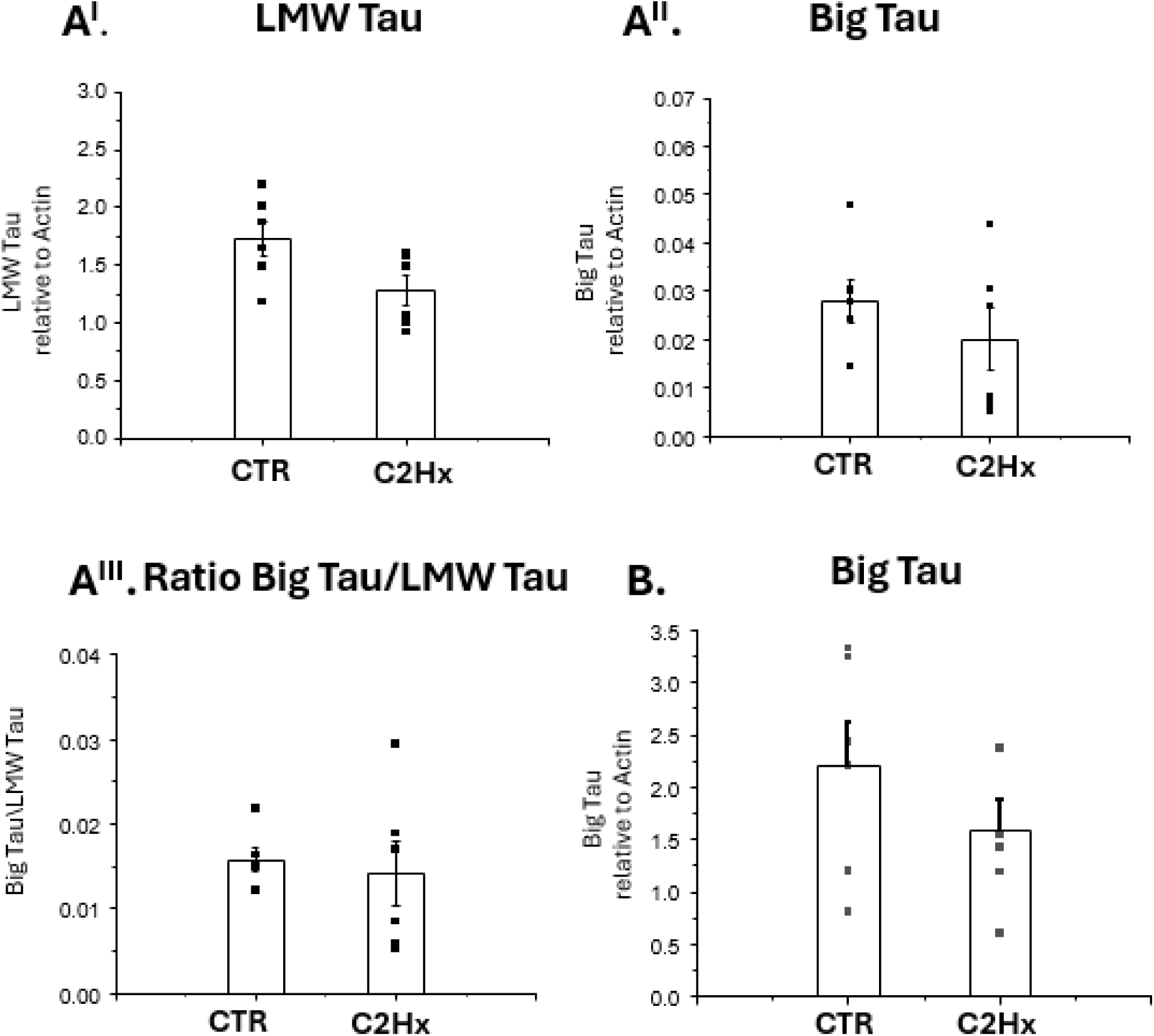
Quantification of Western blot data (A^I^) Quantification of low molecular weight (LMW) tau relative to Actin; (A^II^) Quantification of Big tau relative to Actin; (A^III^) Ratio of Big tau to LMW tau. Panels A^I^–A^III^ represent quantification of the Western blots shown in Figure 2A using the 3′ tau antibody. (B) Quantification of Big tau relative to Actin from Figure 2B, using the Big tau specific antibody shown in Figure 2B. No significant differences were observed between groups (t-test, p>0.05). CTR, control; C2Hx, injury group. Bars represent mean ± SEM, with individual data points shown.

Whereas tau levels showed no significant differences across groups, a significant shift in the tau phosphorylation state was observed following C2Hx. Western blot analysis was performed using the AT8 antibody to detect tau phosphorylation at the Ser202 and Thr205, an epitope widely associated with pathological tau phosphorylation (Rankin et al., 2005), with GAPDH utilized as a loading control to ensure equal protein loading. Figure 4A shows an example of Western blot for control and injured animals. Quantification and statistical analysis (Figure 4B) revealed a significant elevation of p-tau levels in the injury group compared to controls (p=0.03, T-test). These results indicate that although the abundance of tau isoform levels is preserved at four weeks post injury, the C2Hx injury model induces pathological modification of the tau pool below the injury.

**Figure 4.**
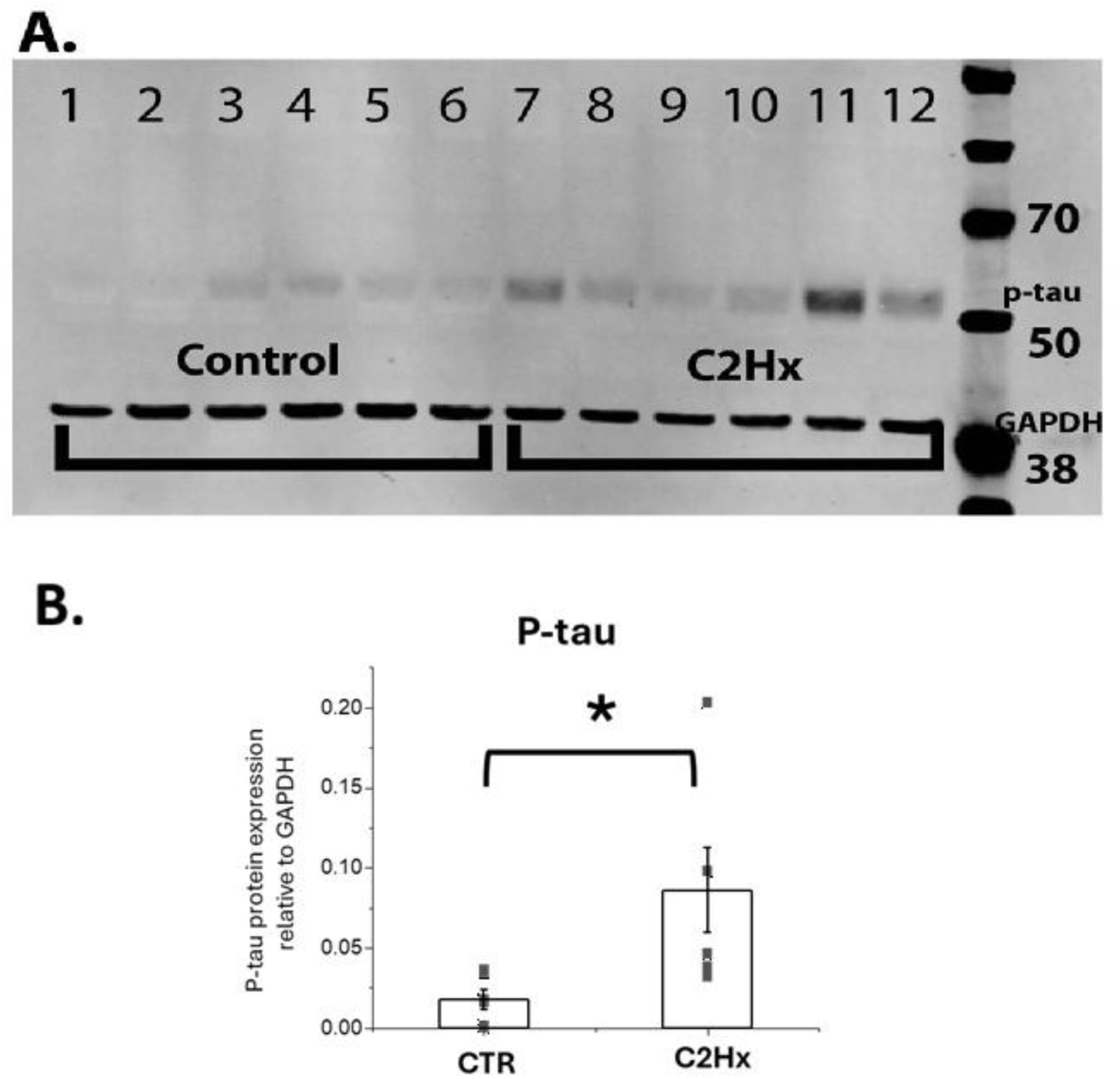
Injury-induced tau phosphorylation at the AT8 epitope. GAPDH was used as a loading control. (A) Representative Western blot. (B) Quantification of phosphorylated tau (p-tau) shows a significant increase in the injured spinal cord compared to controls (p = 0.03). Bars represent mean ± SEM, with individual data points shown. Asterisk (*) marks statistically significant differences between groups (p<0.05).

To determine if the observed increase in tau phosphorylation is associated with changes in microtubule dynamics, the levels of acetylated tubulin (a marker of stable microtubules) and tyrosinated tubulin (a marker of dynamic microtubules) were evaluated. Western blot was performed on the ipsilateral cervical tissue, with Cofilin used as a loading control to ensure consistent protein quantification across samples. Figure 5 shows representative Western blots, while Figure 6 presents the corresponding quantification. Statistical analysis revealed a significant increase in acetylated tubulin levels within the injury compared to sham-operated controls (p=0.001, T-test). In contrast, tyrosinated tubulin levels were significantly higher in the control group, indicating a reduction in microtubule dynamics following C2Hx (p=0.003, T-test).

**Figure 5.**
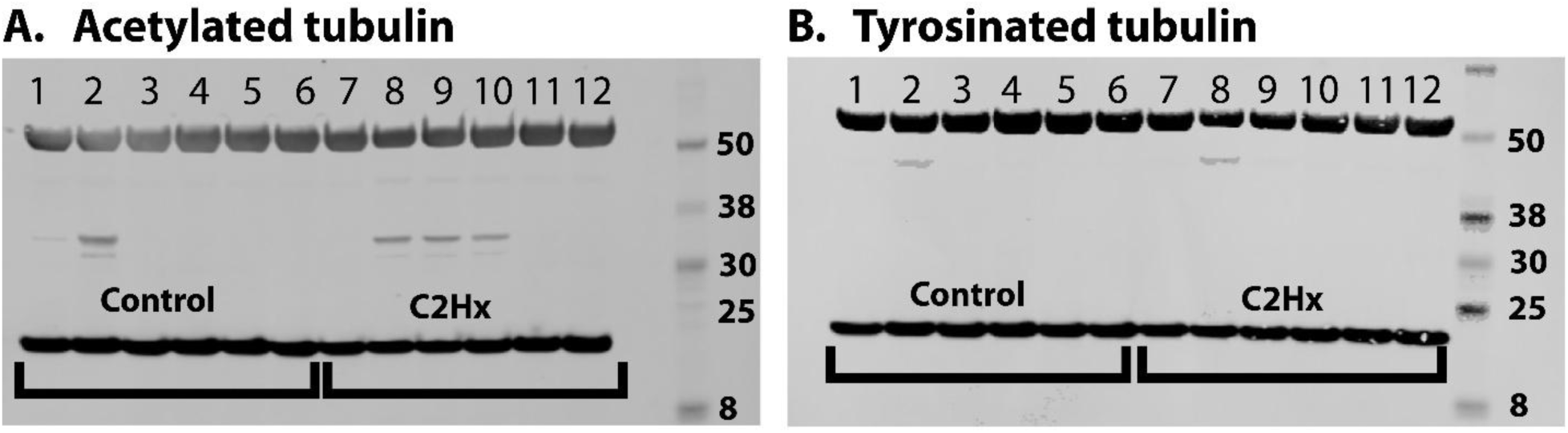
Injury-induced changes in markers of microtubule stability and dynamics. (A) Representative Western blot of acetylated tubulin (50-55 kDa) and (B) tyrosinated tubulin (50-55 kDa) in spinal cord tissue. Cofilin (19-21 kDa) was used as a loading control.

**Figure 6.**
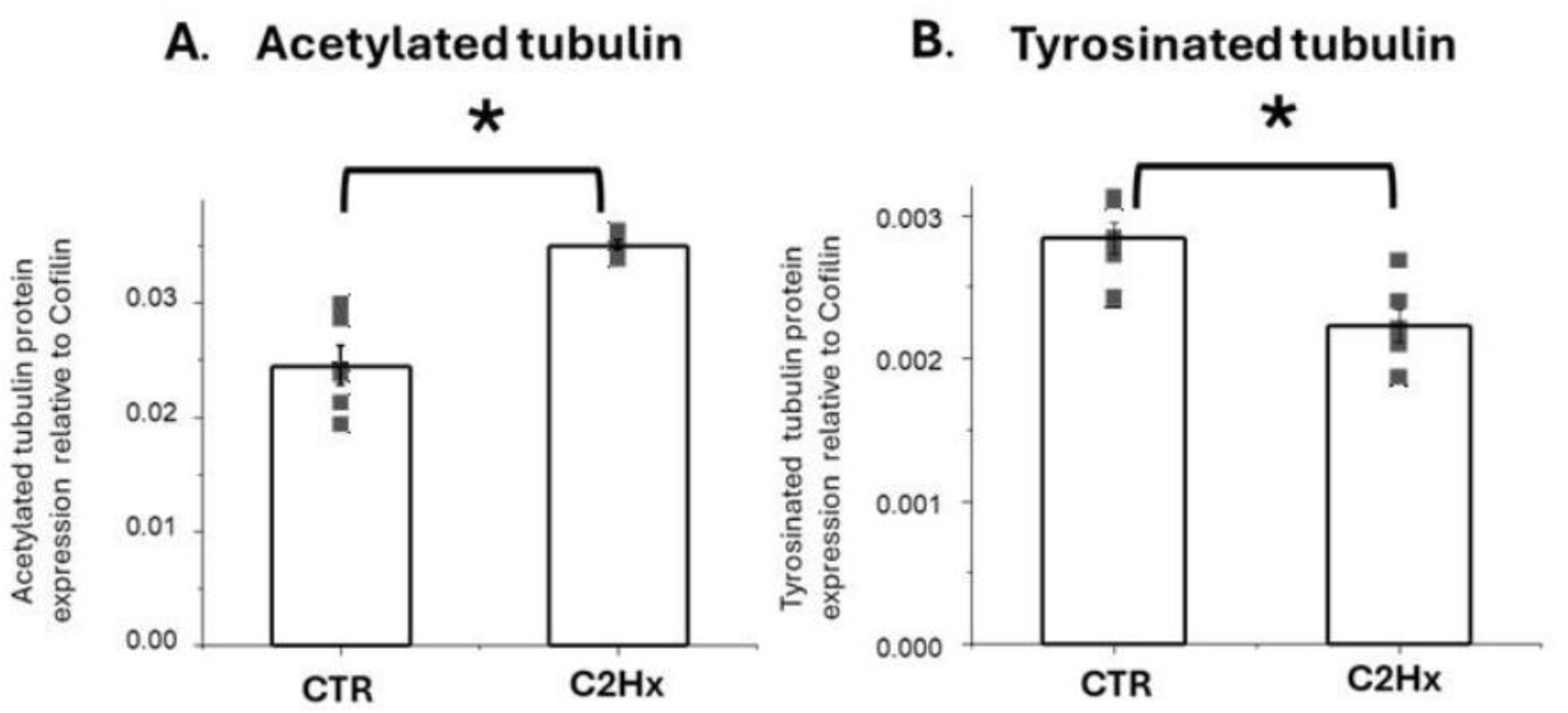
Quantification of acetylated (A) and tyrosinated tubulin (B) normalized to Cofilin. Note significant differences between CTR-control and C2Hx (injury) groups. Bars represent mean ± SEM, with individual data points shown. Asterisk (*) marks statistically significant differences between groups (p<0.05).

### Spatial characterization of tau isoforms and phosphorylation in the injured spinal cord

To map the anatomical distribution of tau pathology associated with p-tau during the early chronic phase of recovery, immunohistochemical analysis was performed on cervical spinal cord sections collected at six weeks post-injury. The study first aimed to identify regions with the highest levels of phosphorylated tau immunoreactivity. As shown in Figure 7A, strong p-tau staining is present at the lesion epicenter, whereas Figure 7B shows p-tau immunoreactivity in tissue caudal to the injury site. P-tau staining was detected in the dorsal horns and multiple white matter regions, including the corticospinal tract (see inset), lateral funiculi, and ventral funiculi.

**Figure 7.**
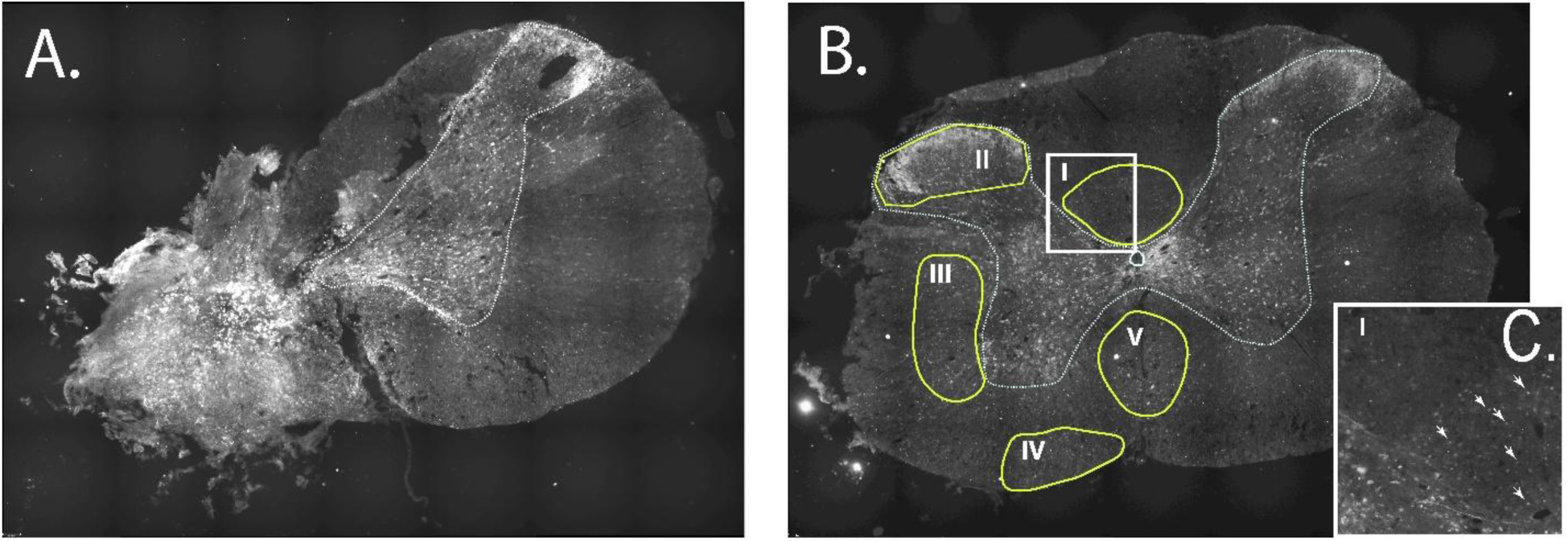
Distribution of phosphorylated tau (p-tau) in the spinal cord at 6 weeks post-injury. (A) Representative p-tau immunostaining at the injury epicenter. (B) Representative p-tau immunostaining at the C3-C4 cervical region. (C) Higher-magnification view of the boxed region in B showing p-tau-positive axons (arrows). I – corticospinal tract; II – dorsal horn; III - lateral funiculus (including bulbospinal pathways); IV – ventrolateral funiculus (including spinothalamic pathways); V-ventral funiculus (including vestibulospinal pathways).

P-tau immunostaining was further examined in the dorsal horn regions caudal and ipsilateral to the lesion. Figure 8 shows representative examples of p-tau and Big tau immunostaining in injured and sham-operated rats. Injured animals showed increased p-tau immunoreactivity in the dorsal horn, whereas Big tau immunoreactivity showed no obvious qualitative difference between groups.

**Figure 8.**
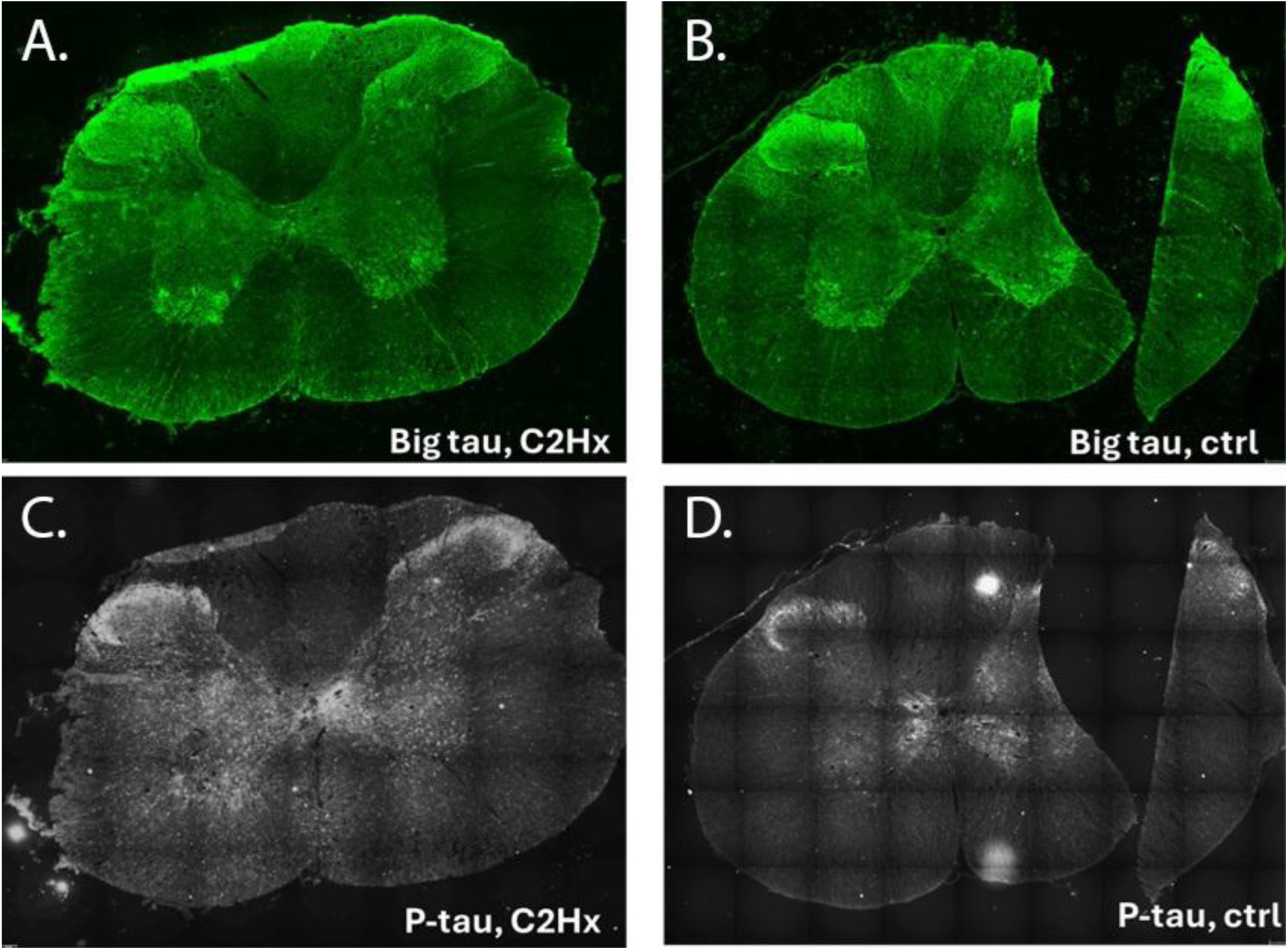
Representative immunohistochemical images of the cervical spinal cord sections (C3-C4) below injury from sham control (ctrl) and C2Hx injured rats at 6 weeks post-injury. All immunohistochemical analyses for these sections were performed at the dorsal horn level. A-B: Immunostaining for Big tau in the (A) injury rat and (B) sham control rat. C-D: Immunostaining for p-tau (AT8) in the (C) injury rat compared to the (D) sham control rat, illustrating a clear increase in phosphorylation specifically within the injured cohort.

Quantification of p-tau and Big tau immunoreactivity was performed in the ipsilateral dorsal horn using sections collected across the C3–C8 cervical spinal cord levels. P-tau immunoreactivity was significantly increased in injured spinal cords compared with sham-operated controls (p = 0.01, t-test, n = 3 per group) (Figure 9A). In contrast, Big tau immunoreactivity was not significantly different between control and injured groups (p = 0.3, t-test, n = 3 per group) (Figure 9B).

**Figure 9.**
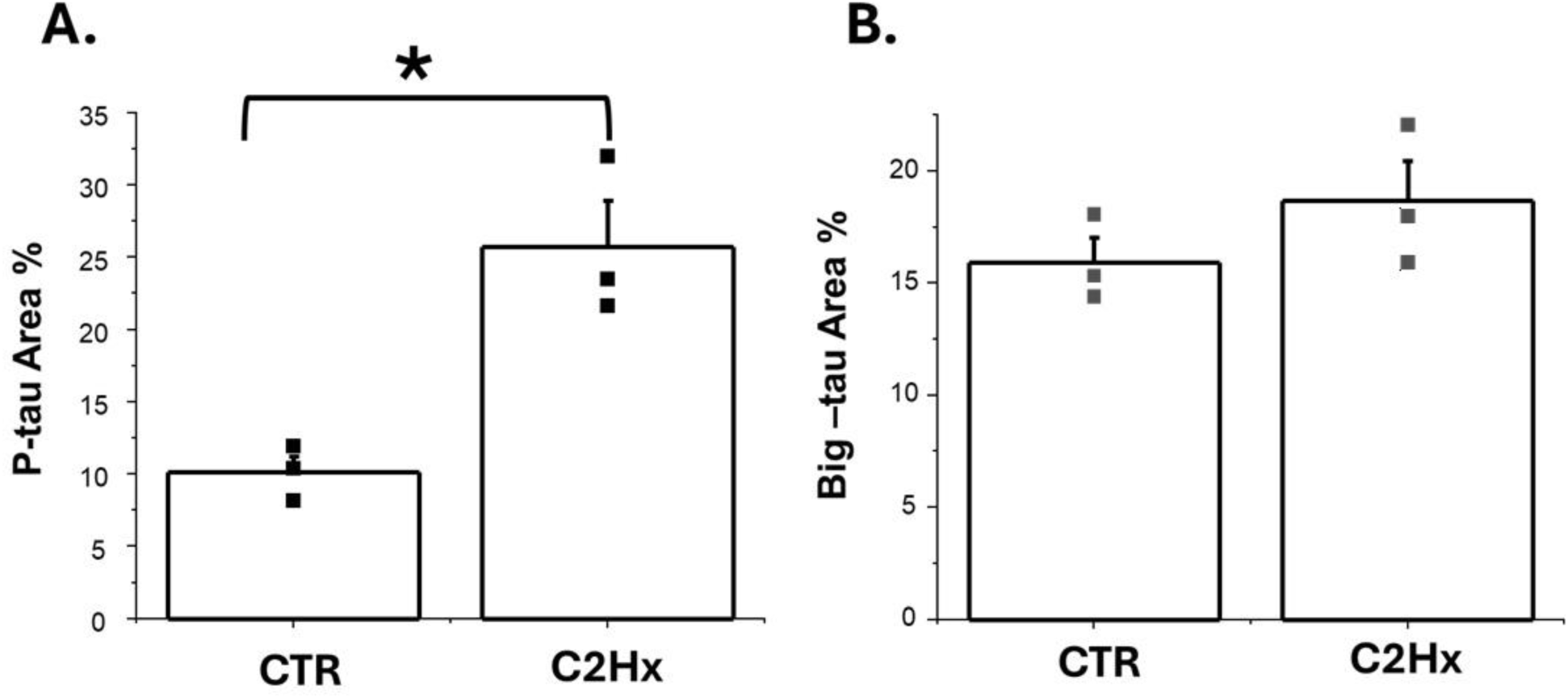
Quantification of phosphorylated (A) and big tau (B) immunoreactivity in the ipsilateral dorsal horn. P-tau-positive area (%) was significantly increased in C2Hx (injured) animals compared to control (CTR) rats (n = 3 per group; p =0.01, T-test). Big tau-positive area (%) was not significantly different between controls and injured rats (n=3, p=0.3, T-test). Bars represent mean ± SEM, with individual data points shown. Asterisk (*) marks statistically significant differences between groups (p<0.05).

To determine whether sensory neurons in the dorsal root ganglia exhibit similar changes in tau phosphorylation, DRGs from cervical levels C6-C7 were analyzed. To address this, DRGs from cervical levels C6–C7 were collected from control (n = 2) and injured (n = 2; lateralized C5 contusion) rats on the ipsilateral side at 8 weeks post-injury (see methods). Double immunostaining was performed using antibodies against Big tau and p-tau. Representative images from control and injured animals are shown in Figures 10 and 11, respectively. Some p-tau–positive neurons exhibited weak Big tau immunoreactivity.

**Figure 10.**
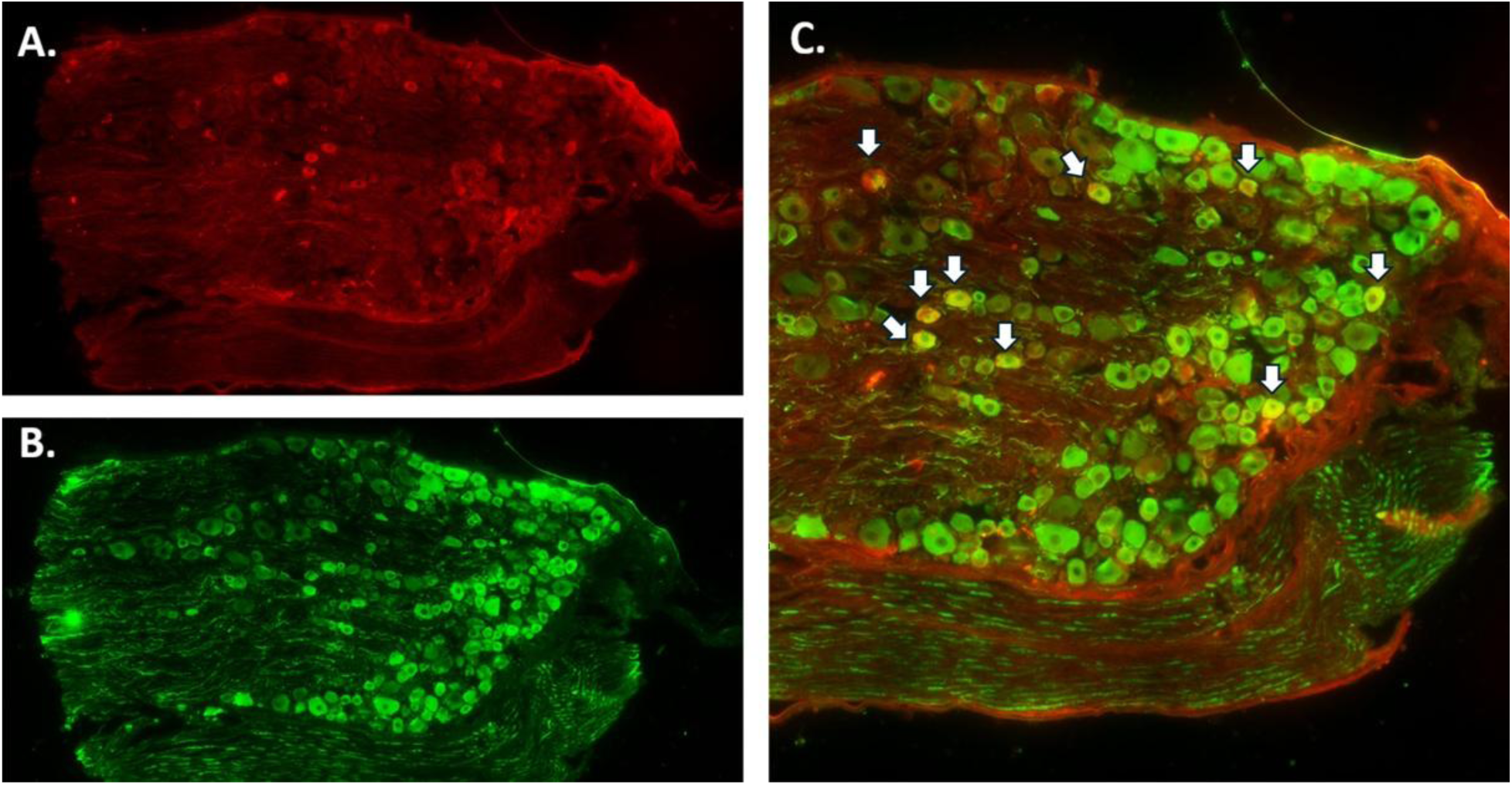
Representative double immunostaining of dorsal root ganglia (DRG) from a control rat 8 weeks post-sham surgery. (A) Phosphorylated tau (p-tau; red) and (B) Big tau (green) immunostaining. (C) Merged image showing co-localization of Big tau and p-tau. White arrows indicate double-labeled neurons.

**Figure 11.**
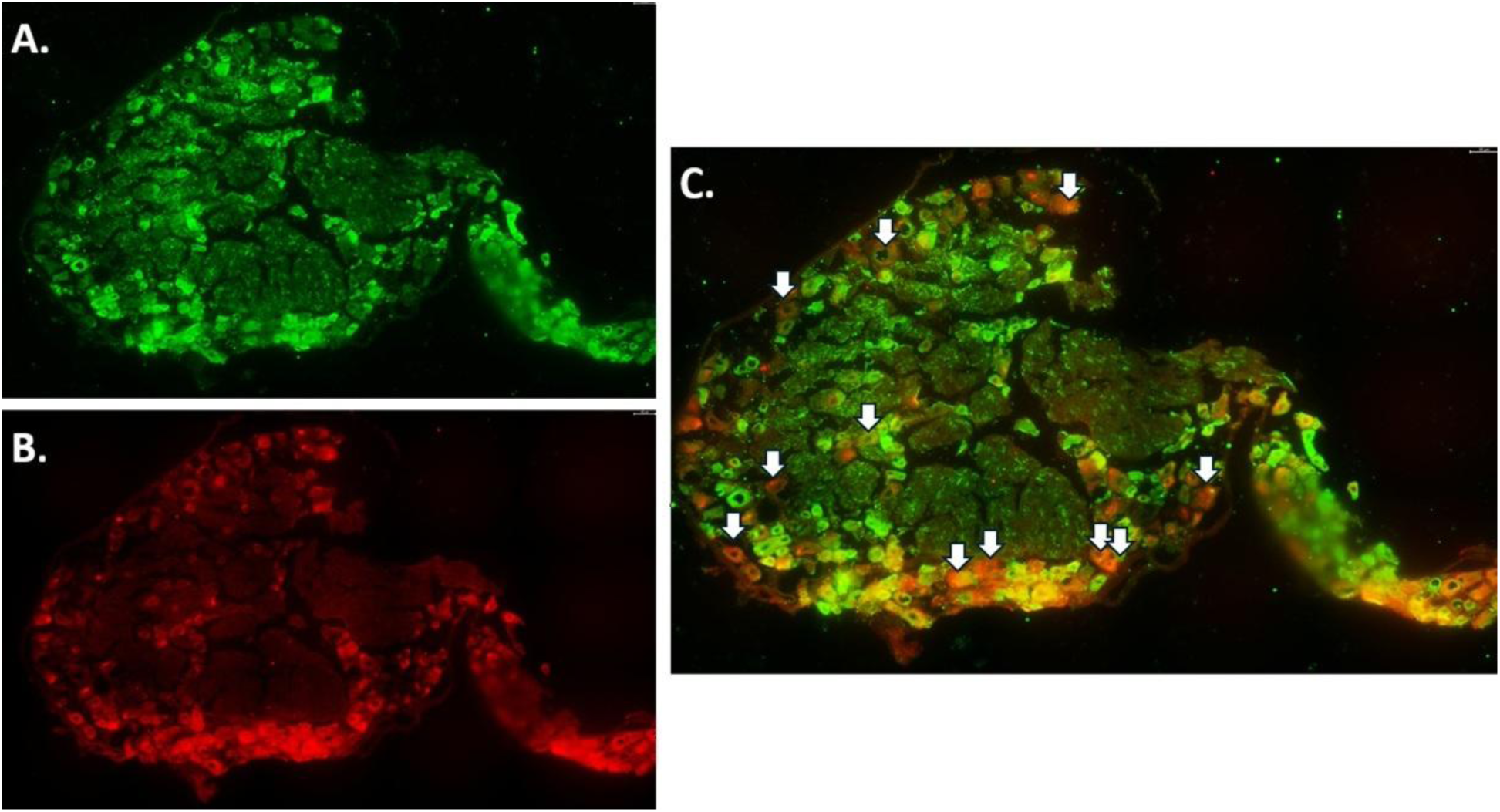
Representative double immunostaining of dorsal root ganglia (DRG) from an injured rat 8 weeks post-injury. (A) Big tau (green) and (B) phosphorylated tau (p-tau; red) immunostaining. (C) Merged image showing co-localization of Big tau and p-tau. White arrows indicate p-tau - positive neurons.

Quantification showed a higher percentage of p-tau-positive neurons in injured DRGs compared with controls (Figure 12). However, because only two animals were available per group, this observation was not subjected to statistical analysis and should be considered preliminary.

**Figure 12.**
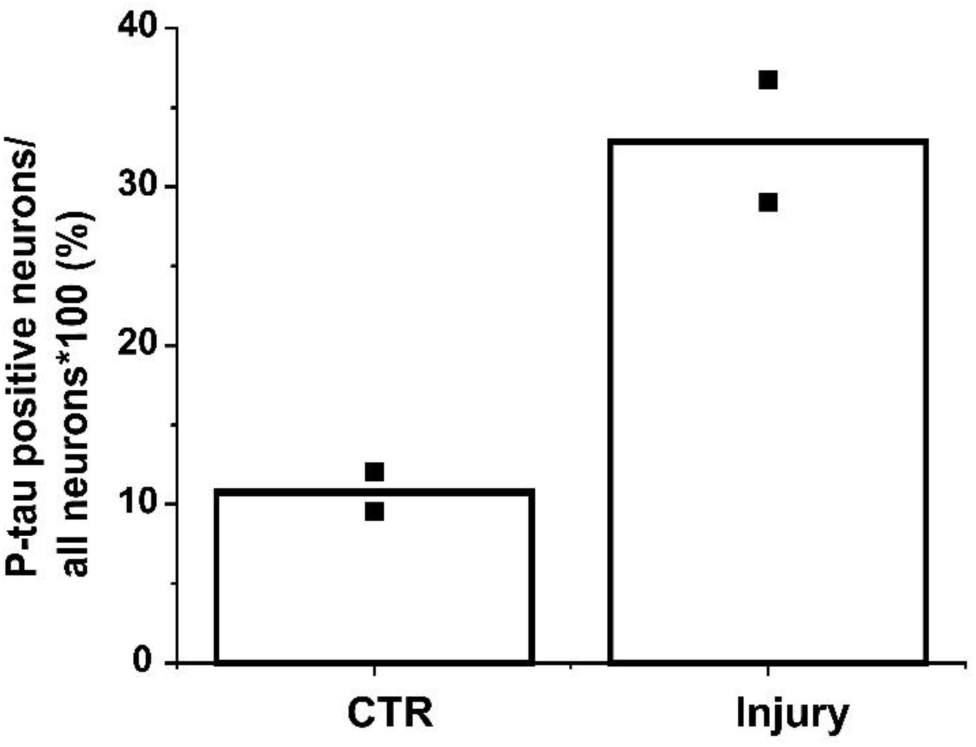
Quantification of p-tau-positive neurons in dorsal root ganglia (DRG) 8 weeks post-injury. The percentage of p-tau-positive neurons relative to all DRG neurons was higher in contused (injured) animals than in control (CTR) animals. Data are shown for two animals per group, with values for each animal averaged across the available sections.

## 4. Discussion

In this study, we investigated alterations in tau signaling following spinal cord injury, with a focus on tau phosphorylation, tau isoforms, and markers of microtubule stability and dynamics. Total tau levels were assessed in control and injured animals. No significant differences in either LMW tau or Big tau levels were found between groups, although a trend toward decreased levels was observed in injured animals (Figures 2–3). We hypothesize that axonal degeneration following a hemisection injury may contribute to reduced tau levels caudal to the lesion. However, such changes are likely to be more pronounced closer to the injury epicenter or within specific regions of the spinal cord. In the present study, the protein analysis was performed using a relatively large segment of spinal cord tissue (C3–C8), which may have masked these localized effects. We also quantified Big tau distribution immunohistochemically in the ipsilateral dorsal horn of control and injured animals (Figure 9B). This analysis did not reveal a significant difference between the two groups, with a similar staining pattern observed in both groups. Immunohistochemical analysis of LMW tau distribution could provide additional information about localized changes following injury. However, we are not aware of an antibody that reliably detects all LMW tau isoforms while excluding Big tau, limiting our ability to perform such an analysis.

A central finding of this study is the significant elevation of phosphorylated tau in injured animals at 4 weeks post-C2Hx (Figure 4). Abnormally increased tau phosphorylation is a well-established feature of tau pathology and is associated with neuronal dysfunction (Avila, 2006). In particular, phosphorylation at the Ser202/Thr205 AT8 epitope examined in the present study has been associated with pathological tau alterations (Rankin et al., 2005). Previous studies using AT8 have reported increased tau phosphorylation both above and below the injury epicenter following contusion injury, peaking at 1 day post-injury and declining by 7 days (Caprelli et al., 2018). In the present study, significantly elevated p-tau levels were observed at 4 weeks post-C2Hx, suggesting that pathological tau phosphorylation may persist beyond the acute phase of spinal cord injury.

In parallel, increased acetylated tubulin and reduced tyrosinated tubulin levels were observed in the injured spinal cord (Figures 5–6). Under physiological conditions, tyrosinated tubulin is associated with newly assembled, dynamic microtubule populations characterized by rapid turnover, whereas acetylated tubulin is enriched in stable, long-lived microtubules (Janke & Magiera, 2020). This balance between dynamic and stable microtubule populations is critical for neuronal plasticity, axonal remodeling, and the formation of new synaptic connections. The accumulation of acetylated tubulin together with reduced tyrosinated tubulin indicates a shift toward a more stable, less dynamic microtubule state following injury.

How might increased tau phosphorylation be associated with a shift toward a more stable microtubule state? The traditional view considers tau primarily as a microtubule-stabilizing protein, with abnormal tau phosphorylation and dissociation from microtubules expected to promote microtubule destabilization (Barbier et al., 2019). This model appears inconsistent with our findings of increased acetylated tubulin and reduced tyrosinated tubulin following C2Hx. However, recent evidence challenges the view of tau as simply a microtubule-stabilizing protein. Tau may instead help regulate microtubule dynamics by maintaining more dynamic regions along axons, thereby enabling axonal microtubules to have long labile/dynamic domains, and changes in tau–microtubule interactions could alter the balance between stable and dynamic microtubules (Baas & Qiang, 2019). In this context, increased tau phosphorylation after C2Hx may contribute to altered microtubule organization, consistent with the concurrent increase in acetylated tubulin and reduction in tyrosinated tubulin observed in the injured spinal cord. Such dysregulation could reduce the cytoskeletal plasticity required for axonal remodeling and regeneration.

Another notable finding of this study is the significant increase in phosphorylated tau within the ipsilateral dorsal horn caudal to the lesion (Figures 7–9). This observation suggests that increased tau phosphorylation may be associated with altered sensory circuitry following spinal cord injury and warrants further investigation into its potential relationship with neuropathic pain (Detloff et al., 2014). However, additional studies are required to test this hypothesis.

An important question arising from this finding is whether the observed p-tau–positive fibers in the dorsal horn originate from primary sensory neurons in the DRG or from intrinsic spinal (propriospinal) sources. Immunohistological staining of DRGs revealed a higher proportion of p-tau-positive neurons in injured animals, suggesting that peripheral sensory neurons may contribute to the increased p-tau signal observed in the dorsal horn following SCI (Figures 10–12). Previous studies have shown that Big tau is predominantly expressed in adult DRG neurons (Jin et al., 2023) and have suggested that its unique structure, which includes the long 4a exon, is less prone to pathological aggregation (Fischer & Baas, 2020). In the spinal cord, the increase in p-tau detected by Western blot with the AT8 antibody occurred within the molecular weight range corresponding to LMW tau, suggesting that the injury-associated increase in tau phosphorylation predominantly involved LMW tau isoforms (Figure 4).

Our Western blot analysis of DRG tissue revealed low levels of LMW tau-immunoreactive bands corresponding in molecular weight to the prominent LMW tau bands observed in spinal cord tissue on the same blot (Figure 13). These findings indicate that LMW tau isoforms are present in DRG tissue and raise the possibility that LMW tau may contribute to the increased p-tau observed in DRG neurons following injury. However, our data do not directly identify which tau isoform is phosphorylated in DRG neurons, and further studies are required to address this question.

**Figure 13.**
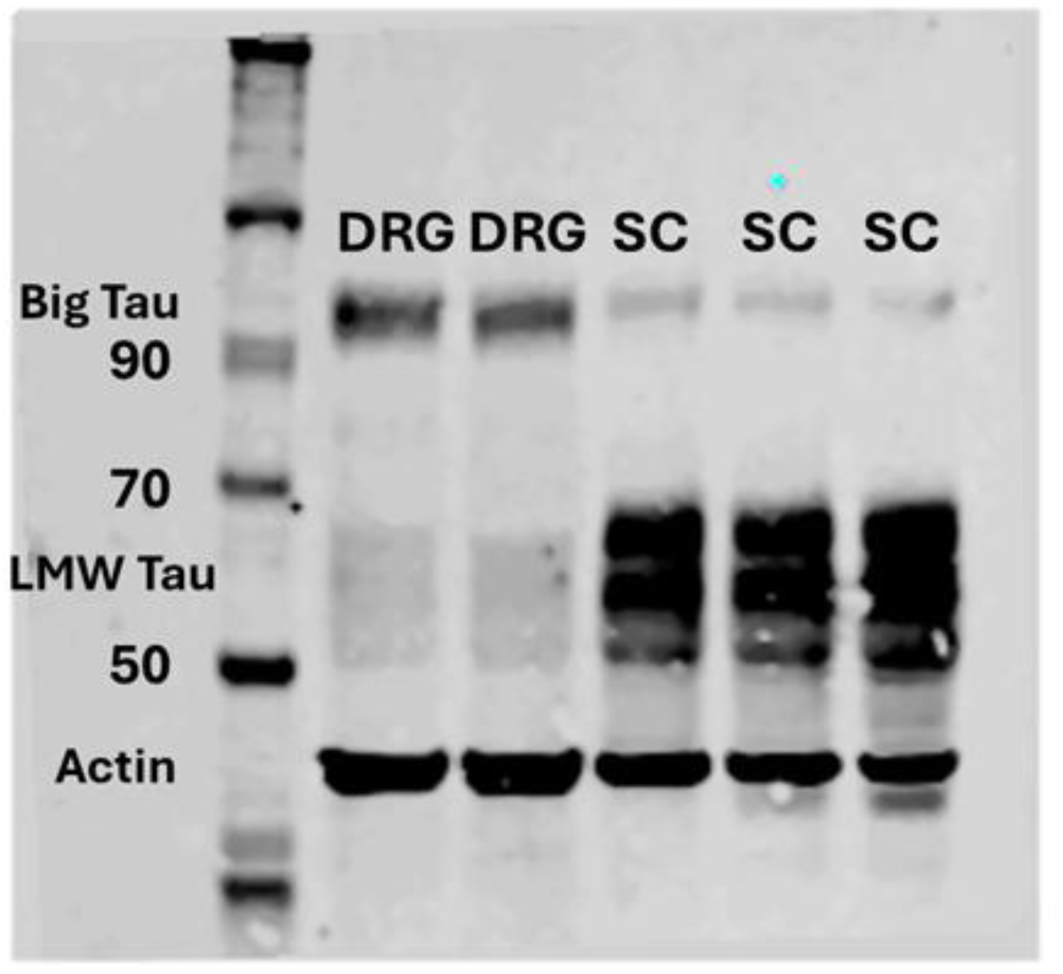
Tau isoform expression in dorsal root ganglia (DRG) and spinal cord (SC). Representative Western blot probed with a 3′ tau antibody showing low molecular weight (LMW) tau (∼50–65 kDa) and Big tau (∼90–110 kDa) in DRG and SC tissue. Spinal cord samples show prominent LMW tau bands with relatively weak Big tau signal, whereas DRG samples show a prominent Big tau band together with weaker bands corresponding in molecular weight to LMW tau species observed in spinal cord. Actin was used as a loading control.

Overall, these findings suggest that persistent alterations in tau phosphorylation and microtubule organization following spinal cord injury may influence plasticity within spared neural circuits and identify tau–microtubule regulation as a potential target for therapeutic intervention.

Pharmacological approaches aimed at reducing pathological tau phosphorylation have been explored in other models (Hung et al., 2005), raising the possibility that modulation of tau signaling may also have therapeutic relevance following SCI. Given the changes observed in both spinal cord and DRG, future studies should determine whether targeting tau phosphorylation can influence not only motor recovery but also sensory dysfunction, including neuropathic pain.

## 5. Summary

The findings of this study suggest that spinal cord injury is associated with persistent dysregulation of tau phosphorylation in both the spinal cord and DRG, together with altered microtubule organization in the injured spinal cord. These changes may influence the capacity of injured neural circuits to undergo adaptive remodeling, although the relationship between tau phosphorylation, microtubule organization, and functional recovery remains to be determined.

## Credit authorship contribution statement

N.B.: Investigation, Data curation, Formal analysis, Visualization, Writing, review & editing. S.C.: Investigation, Methodology, Resources, Validation, Technical support. T.B.: Conceptualization, Methodology, Investigation, Formal analysis, Data curation, Writing, review & editing, Supervision, Funding acquisition.

## Acknowledgments

This work was supported in part by the National Institutes of Health (NIH) under award R21NS131962 and by Grant #3178 from the Paralyzed Veterans of America Research Foundation. The authors thank Dr. Megan Detloff for generously providing the dorsal root ganglion tissue samples used in this study. The authors also thank Dr. Itzhak Fischer for generously providing the 3′ tau and big tau antibodies, as well as for his helpful discussions and critical review of the manuscript.

## Data availability

Data will be made available on request.

